# Pre-anthesis light signaling of sheathed embryonic barley inflorescences defines floral fate

**DOI:** 10.64898/2025.12.01.691580

**Authors:** Sai Thejas Babanna, Yongyu Huang, Andreas P.M. Weber, Thorsten Schnurbusch

## Abstract

The pre-anthesis embryonic inflorescence greening (PEIG) is a distinctive developmental feature in the members of the Triticeae such as barley, wheat and rye. In barley (*Hordeum vulgare* L.), floral survival and fertility are major determinants of grain yield, yet the physiological processes supporting early inflorescence development remain poorly understood. Here, we investigated PEIG, a light-dependent chlorophyll accumulation occurring while immature inflorescences are still enclosed by leaf sheaths, and its role in overall inflorescence development. Using chlorophyll autofluorescence imaging and chlorophyll quantification, we found that PEIG in barley initiates at a surprisingly early developmental stage, when developing spikes are still enclosed by leaf sheaths. PEIG first appears in the central rachis and then progressively spreads to the spikelet primordia and other floral organs. Using a non-destructive dark treatment method designed for preventing light exposure to developing inflorescences, we further demonstrated that the inhibition of PEIG does not affect floral initiation but significantly reduces floral survival and pollen viability, particularly in the tip region of the inflorescence, indicating a critical role of light mediated PEIG in floral fate. Finally, analysis of natural variation for PEIG revealed a strong positive correlation between PEIG and floral survival. Our findings establish PEIG as an underappreciated hidden trait that supports floral viability and reproductive success, with implications to enhance grain yield potential in cereals.

## 1. Introduction

Light is a central regulator of plastid differentiation, photosynthetic gene expression, and chlorophyll biosynthesis (Cackett et al., 2022; Pogson et al., 2015). It also strongly affects reproductive development, with light intensity, photoperiod, and light quality influencing overall floral fertility (González et al., 2003, 2005; Parrado, Savin, et al., 2025; Parrado, Slafer, et al., 2025). Several shading experiments clearly demonstrate that reduced irradiance during reproductive development can significantly diminish grain yield (Dong et al., 2019; Slafer et al., 1994), largely due to insufficient assimilate supply and impaired source-sink balance (Masanori et al., 2001; Yang et al., 2020). By regulating chloroplast development and reproductive organ formation, light directly influences the photosynthetic activity of the inflorescence, which supports grain development.

In cereals, fully developed inflorescence organs including awns, lemma, palea and glumes containing functional chloroplasts have been shown to significantly contribute to the final grain yield, especially in the later stages of development when the leaves start to senesce (Gebbing & Schnyder, 2001; Imaizumi et al., 1990; McKenzie, 2011; Sanchez-Bragado et al., 2016). These chlorophyllous reproductive structures supply assimilates to developing grains and other sink tissues within the spike (AuBuchon-Elder et al., 2020; X. Li et al., 2006). In Triticeae species, such as barley (*Hordeum vulgare* L.), wheat (*Triticum aestivum* L.) and rye (*Secale cereale* L.), chlorophyll accumulation in the inflorescence begins at an early developmental stage, even before anthesis, while the inflorescence is still enclosed by multiple layers of leaf sheaths (Huang et al., 2023; Shanmugaraj et al., 2023). The pre-anthesis embryonic inflorescence greening (PEIG) is a characteristic feature of Triticeae species, which occurs even before emergence (Huang et al., 2023).

Inflorescence development is a major determinant of reproductive success in cereals. Barley has a spike-type inflorescence in which spikelets, the basic floral grain-bearing structures, are arranged in an alternate-distichous pattern along the central axis (the rachis). Inflorescence architecture, including both the total number of initiated spikelets and the proportion that ultimately survive to set grain, strongly influences yield potential (Arisnabarreta & Miralles, 2008; Kamal, et al., 2022; Shanmugaraj et al., 2023). Barley, possessing an indeterminate inflorescence structure, initiates a large number of spikelet primordia during the early reproductive development, while the end of the spikelet primordia initiation along the rachis marks the maximum yield potential stage (MYP) (Koppolu & Schnurbusch, 2019; Thirulogachandar & Schnurbusch, 2021). However, a substantial fraction of these primordia degenerate before anthesis, particularly in the apical region, resulting in the pre-anthesis tip degeneration (PTD) (Huang et al., 2023; Shanmugaraj et al., 2023). Both MYP and PTD differ among barley genotypes and ultimately final grain number (Kamal, et al., 2022; Thirulogachandar & Schnurbusch, 2021). Therefore, understanding the physiological and developmental factors influencing spike PTD may thus ultimately help improve grain yield.

Recent research, using genetic and transcriptomic approaches, has highlighted the critical role of early chlorophyll accumulation in developing barley inflorescences (Huang et al., 2023; Shanmugaraj et al., 2023). For example, barley CMF4 (HvCMF4), a CCT-motif family protein, was identified as a key regulator orchestrating chloroplast development, light signaling, and vascular differentiation in the embryonic inflorescence. Knockout of *HvCMF4* reduced the overall spike chlorophyll accumulation, particularly in the apical region of the developing inflorescence, thereby impairing spike development. This phenotype was further recapitulated by applying photosynthesis inhibitors (norflurazon and lincomycin) to the plant, underscoring the important role of PEIG in supporting proper inflorescence development (Huang et al., 2023). Complementary transcriptomic analyses in barley showed that developing spikes, even though being covered by several layers of leaf sheaths and exposed to dark conditions, express numerous chloroplast biogenesis, chlorophyll biosynthesis, and photosynthesis-related genes long before emergence (Shanmugaraj et al., 2023). Together, these findings highlight PEIG as an overlooked yet potentially important developmental cue, suggesting that the timing and spatial patterning of greening within developing spikes may be functionally linked to spikelet survival, reproductive development, and overall reproductive success.

Although light is known to influence adult inflorescence and grain development after heading, it is unclear whether embryonic spikes can perceive light, particularly for initiating PEIG while still being enclosed within leaf sheaths. The extent to which PEIG depends on light, and whether limiting light affects spikelet survival or reproductive development, remains unknown. These knowledge gaps highlight the need for a mechanistic understanding of PEIG and its developmental function. Despite recent discoveries that barley spikes initiate chlorophyll biosynthesis well before head emergence (Huang et al., 2023; Shanmugaraj et al., 2023), the timing, spatial progression, and physiological relevance of this process have not been systematically examined. Although photosynthesis inhibitors have been applied to whole plants in attempts to inhibit PEIG (Huang et al., 2023), no studies have experimentally disrupted PEIG without affecting overall plant growth to directly test its role in spikelet initiation, survival, or fertility, nor has natural variation in PEIG been linked to genotypic differences in spikelet survival. To address these gaps, we combined high-resolution imaging, chlorophyll quantification, a non-destructive dark-treatment approach, and a diverse barley panel to characterize PEIG and determine its developmental significance. Our findings reveal that early greening is a critical physiological cue supporting meristem viability and reproductive success, thereby establishing PEIG as a previously unrecognized target for improving spike fertility and yield potential in barley.

## 2. Materials and Methods

### 2.1 Plant material and growth conditions

#### Green house experiments

Greenhouse experiments (photoperiod: 16 hours/8 hours, light/dark; temperature: 20°/16°C, light/dark) were conducted at the Leibniz Institute of Plant Genetics and Crop Plant Research (IPK) in between 2023-2025. For both phenotyping and dissection of immature spikes experiments, two-rowed barley cv. Bowman (hereafter Bowman) grains were germinated in a 96-well planting tray for 2 weeks, vernalized at 4°C for 2 weeks, acclimatized at 15°C for a week, and lastly potted into 9cm pots until maturity or until sample was collected.

#### Field experiments

Both phenotyping and dissection experiments were done in field condition at IPK in 2025. The field containing natural loamy soil was manually watered and fertilized. Twenty genotypes of six rowed spring barleys were selected from the ∼ 350-barley diversity panel coming from Federal Ex-situ Gene Bank hosted at IPK. (Huang et al., 2023; Kamal et al., 2022) (Table S1). Barley grains were germinated in jiffy pots in greenhouse conditions, two weeks old seedlings along with the jiffy pots were transferred to the semi-field conditions with uniform distance of 10-12 cm between each plant and a distance of 20-25 cm was maintained between each row. Spike samples were collected by dissection upon reaching the right stage and for phenotyping, plants were grown until maturity.

### 2.2 Phenotyping

All phenotypic data from greenhouse-grown plants were collected from the main culm. Final spikelet number was recorded between heading and the grain-filling stage, while grain number, dry biomass, and tiller number were measured at harvest. For plants grown under semi-field conditions, the first three spikes were averaged to determine spikelet number between heading and grain filling, and the same spikes were subsequently used to assess grain number after harvest. Spikelet survival and spike fertility were calculated as follows:

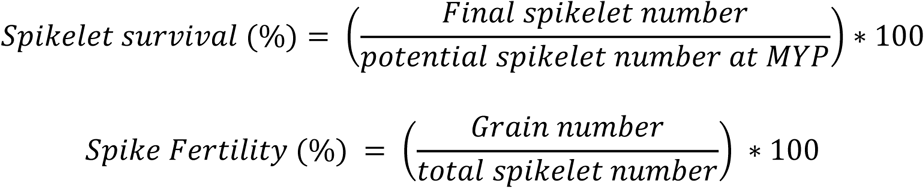

### 2.3 Treatment and experimental design

For the dark treatment (DT) experiments, three treatment groups (DT_W2-10_, DT_W2-5.5_, and DT_W4.5-10_) were included along with a control. To prevent light from reaching the developing inflorescence, we developed a method using an aluminum foil strip to directionally shield the targeted region without interfering with overall plant growth or photosynthesis (Fig. S1 & 2b). Prior to applying the treatment, plants at comparable developmental stages were dissected to determine the precise position of the internode bearing the developing inflorescence (Fig. S1a). Based on this position, an aluminum foil strip (1–1.5 cm wide) was gently wrapped around the leaf sheaths at the first internode on the main culm (Fig. S1b). The strip was wrapped in a circular manner to ensure complete coverage of the developing inflorescence and was positioned to cover approximately 2-3 cm above and below the spike position depending upon the stage and size of the inflorescence (Fig S1c). Further, untreated reference plants of the same developmental stage were monitored to track the exact position of the inflorescence, and the foil wrap was adjusted as needed to maintain full coverage throughout development. The aluminum wrap was applied loosely to avoid any physical restriction to the developing inflorescence. This method allowed passive gas exchange and did not damage surrounding tissues, thereby preventing any direct or indirect effects on overall plant development.

#### Temperature measurements

To measure the local temperature beneath the aluminum-foil cover and around the developing inflorescence, the position of the immature spike at ∼W4.5 was first identified as mentioned earlier (Fig. S2a). A digital thermometer with a long probe was then positioned so that the probe tip rested exactly where the spike was located (Fig. S2b), after which the aluminum foil was wrapped around the leaf sheaths together with the thermometer probe (Fig. S2c & d). This allowed measurement of the ambient temperature beneath the foil cover (Fig. S2 and Fig. 3a). Temperature readings were recorded every 2 hours throughout the day while the plant remained at the W4.5 stage. Three biological replicates with aluminum-foil covers were monitored, and an additional three uncovered control replicates were used to measure ambient temperature conditions.

### 2.4 Chlorophyll measurements

For chlorophyll measurements the IMs at W4.5-5 and W6 stages were collected by manually dissecting the main culm and further the tissue was frozen immediately using liquid nitrogen in a 2-ml Eppendorf tube. For each biological replication ∼15 spikes (W4.5 - 5) and ∼10 (W6) were pooled. Sample collection was done between 11:00 a.m. and 2:00 p.m. of a day. Chlorophyll concentration was measured according to (Porra et al., 1989). After grinding the tissue samples into fine powder, 1ml of methanol was added followed by centrifugation at 1300rpm for two minutes. Supernatant was collected in clean Eppendorf tubes labelled as set one. In the remaining supernatant, 1ml of methanol was added again, followed by centrifugation and supernatant collection in the second set of Eppendorf tubes. The final sample was prepared by taking 600μl of supernatant from each set, making the final volume 1200μl. From the final volume, 800μl was used for the downstream spectrophotometer measurements. The measurements were taken at two wavelengths, i.e., 652nm and 665.2nm. Final chlorophyll (a+b), chlorophyll a and chlorophyll b concentration was calculated as follows:

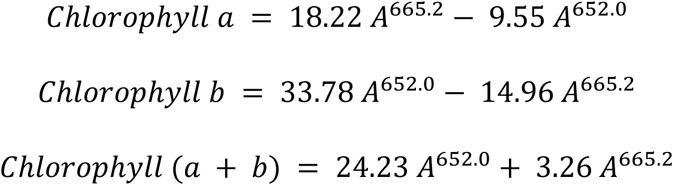

### 2.5 Microscopy

Chlorophyll autofluorescence in IMs was examined using confocal laser scanning microscopy (CLSM). Immature inflorescences at developmental stages W4.5-W8.5 were dissected and embedded in 7% agarose within a flat-bottom mold. After solidification, agarose blocks were removed and sectioned into 80–90 µm slices using a vibrating blade microtome (Leica VT1000 S). For earlier stages (W2–W3.5), IMs were mounted directly on microscope slides in water and covered with a coverslip. Autofluorescence was captured on an LSM780 CLSM system (Carl Zeiss MicroImaging) using a 640-nm laser line and emission from 650 to 720 nm.

### 2.6 Pollen viability

Pollen viability was evaluated using the Amphasys® impedance flow cytometer method. Spikes from all DT and control group plants grown in the greenhouse were collected when anthers were near anthesis. Outer floral organs (lemma and palea) were carefully removed to expose mature, yellow, swollen anthers close to dehiscence. For each replicate, three anthers from a single spikelet were collected prior to opening and transferred into Eppendorf tubes containing Amphasys® buffer 11. Samples were filtered through 100 µm filter heads and analyzed by IFC. Pollen passed through a 120 µm chip, where an electric field was applied, and impedance phase shifts were recorded. Viable and dead pollen were distinguished using a gating threshold calibrated with dead reference samples prepared by boiling pollen at 100 °C for 1 h. The resulting pollen viability data coming from 5-8 biological replicates were subsequently used for downstream analyses.

### 2.7 Statistical analysis

All the data analysis and figure generation in this study was done using RStudio Software. Packages such as ggplot2, dplyr, and ggpubr were used for all the data processing and analysis. Unpaired Student’s t-test analysis for all comparisons was performed to check the significance of the data. The resulting p-value is represented by: ‘*’ indicates p-value ≤ 0.05; ‘**’ ≤ 0.01; ‘***’ ≤ 0.001; ‘****’ ≤ 0.0001; ‘ns’ – not significant. Pearson’s correlation was performed to assess linear relationships between traits.

## 3. Results

### 3.1 Initiation of greening in the immature inflorescences

To determine the onset and spatial progression of chlorophyll accumulation during the development of immature barley inflorescences, which are surrounded by several layers of leaf sheaths, we examined embryonic spikes at defined stages according to the Waddington scale (Waddington et al., 1983). Observations for PEIG started at Waddington stage 1.5 (W1.5), a critical point at which the shoot apical meristem transits from the vegetative to the reproductive phase. At W1.5, a clear chlorophyll autofluorescence signal was already detectable within the immature spikes, indicating the initiation of chlorophyll biosynthesis in the developing rachis area (Fig. 1a). The intensity of the auto fluorescence signal increased as development progressed (Fig. 1a-f), showing even stronger signal by stage W4.5 (Fig. 1d). Spatially, the earliest appearance of chlorophyll was localized mainly to the central portion of the developing meristem, corresponding to the developing rachis region (Fig. 1a-e). As development proceeded, chlorophyll accumulation expanded to the spikelet primordia and subsequently the emerging floral organs, such as the awns, lemma and palea (Fig. 1d-f). Within the rachis, a distinct acropetal gradient of chlorophyll distribution was persistently observed; the basal region of the immature spike exhibited the strongest auto fluorescence signal, which gradually reduced toward the distal tip, eventually disappearing altogether (Fig. 1d & e). Further quantitative analysis of total chlorophyll content in whole spikes supported these imaging results. A steady increase in chlorophyll levels was detected between stages W4.5 and W5.5 (Fig. 2a), corroborating the visual evidence of enhanced greening and suggesting that chlorophyll biosynthesis accelerates during this developmental window. These results demonstrate that chlorophyll accumulation begins at earliest stage of inflorescence development and intensifies as inflorescence development progresses.

**Figure 1:**
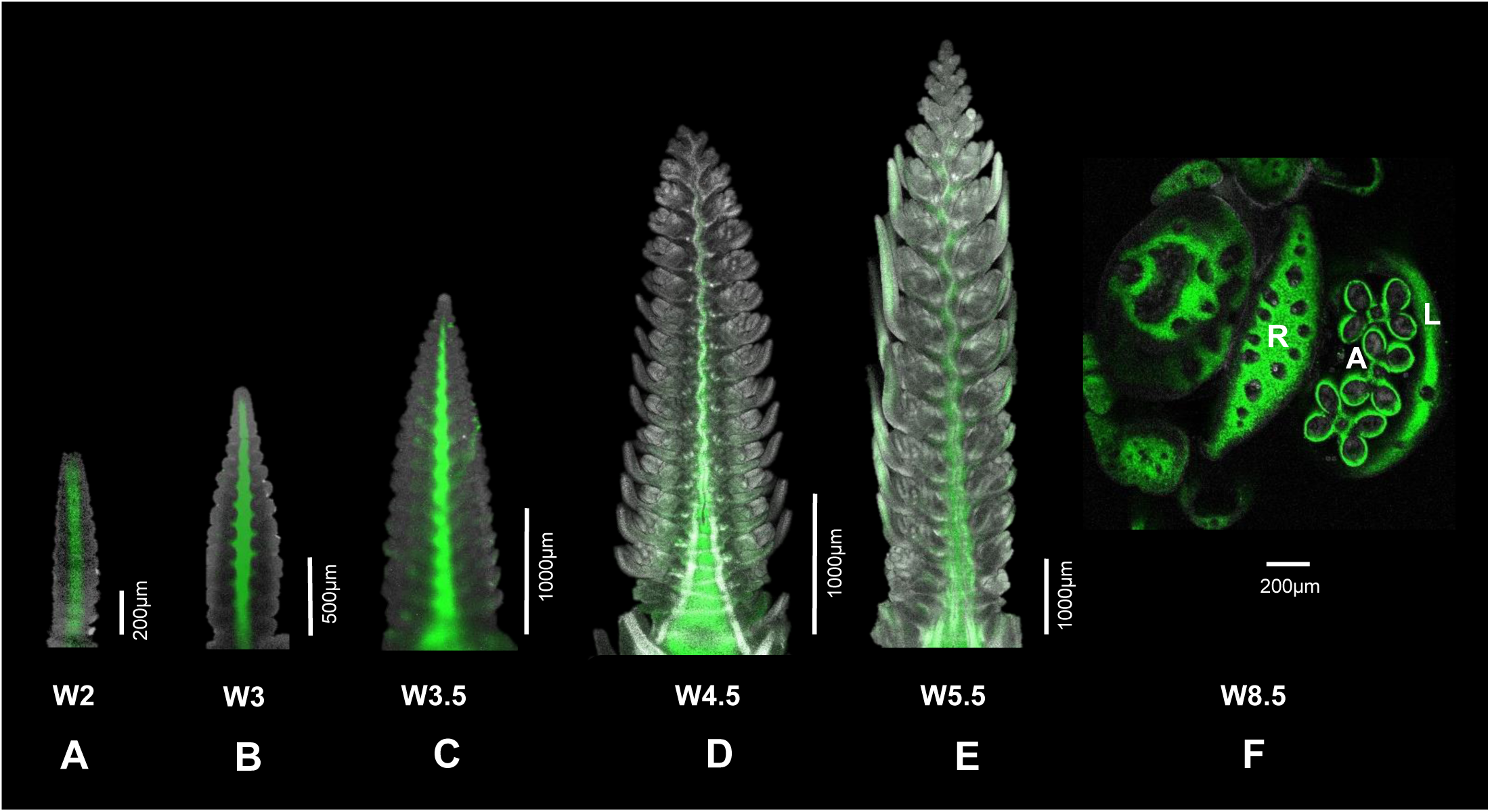
(A-F) Distribution of chlorophyll in the developing inflorescence, visualized by auto chlorophyll fluorescence at different Waddington developmental stages. R - rachis; A - anther; L - lemma.

**Figure 2:**
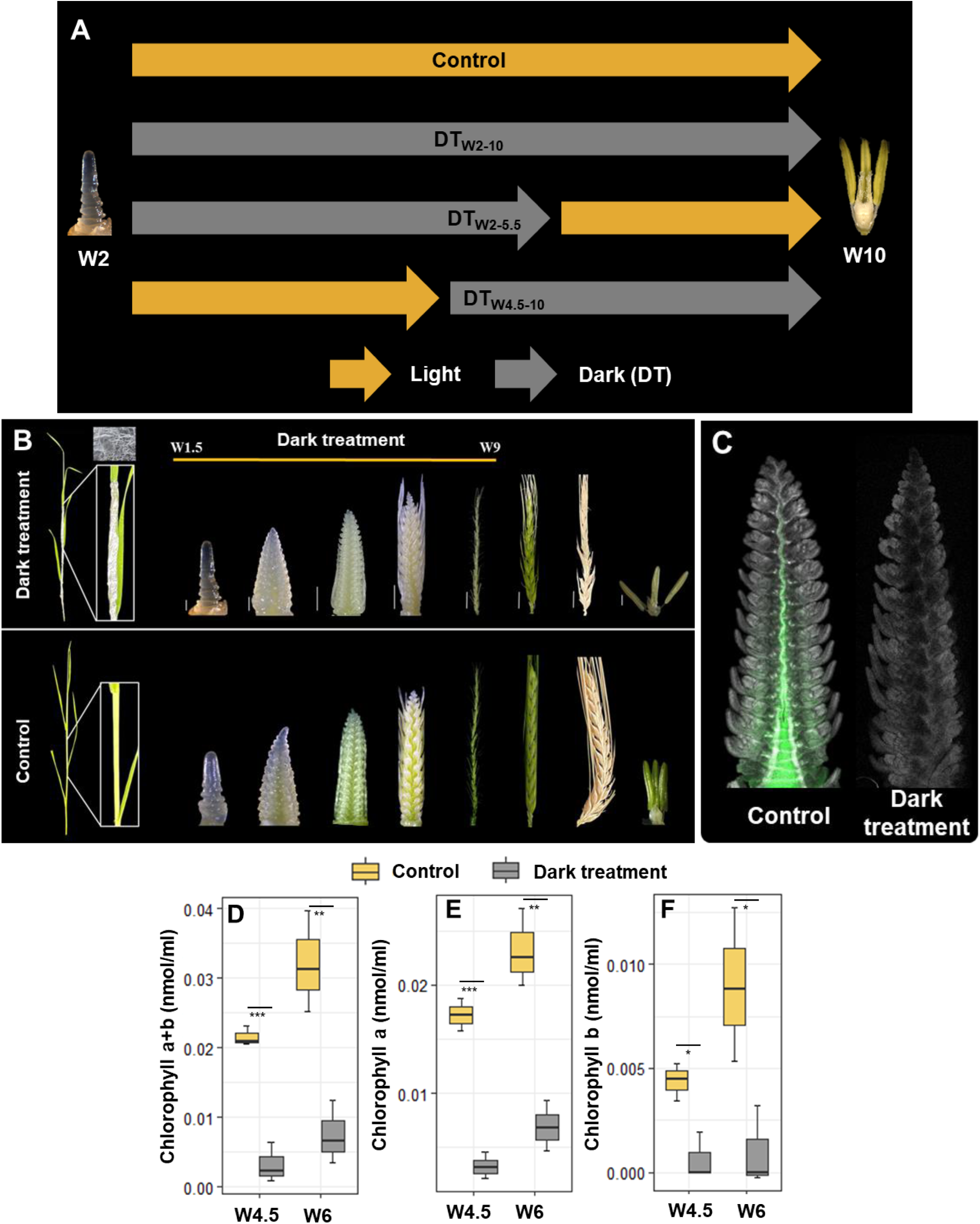
Effects of dark treatment on developing inflorescences. (A) Schematic representation of the treatment groups and their respective treatment phase. (B) Images of the developing inflorescences from dark treated and control treatments at different developmental stage between W1.5-W9+. (C) Chlorophyll autofluorescence in control (left) and dark-treated (right) inflorescence meristems at W4.5. Chlorophyll concentration in immature spike tissues, (D) chlorophyll a+b, (E) chlorophyll a, (F) chlorophyll b

**Figure 3:**
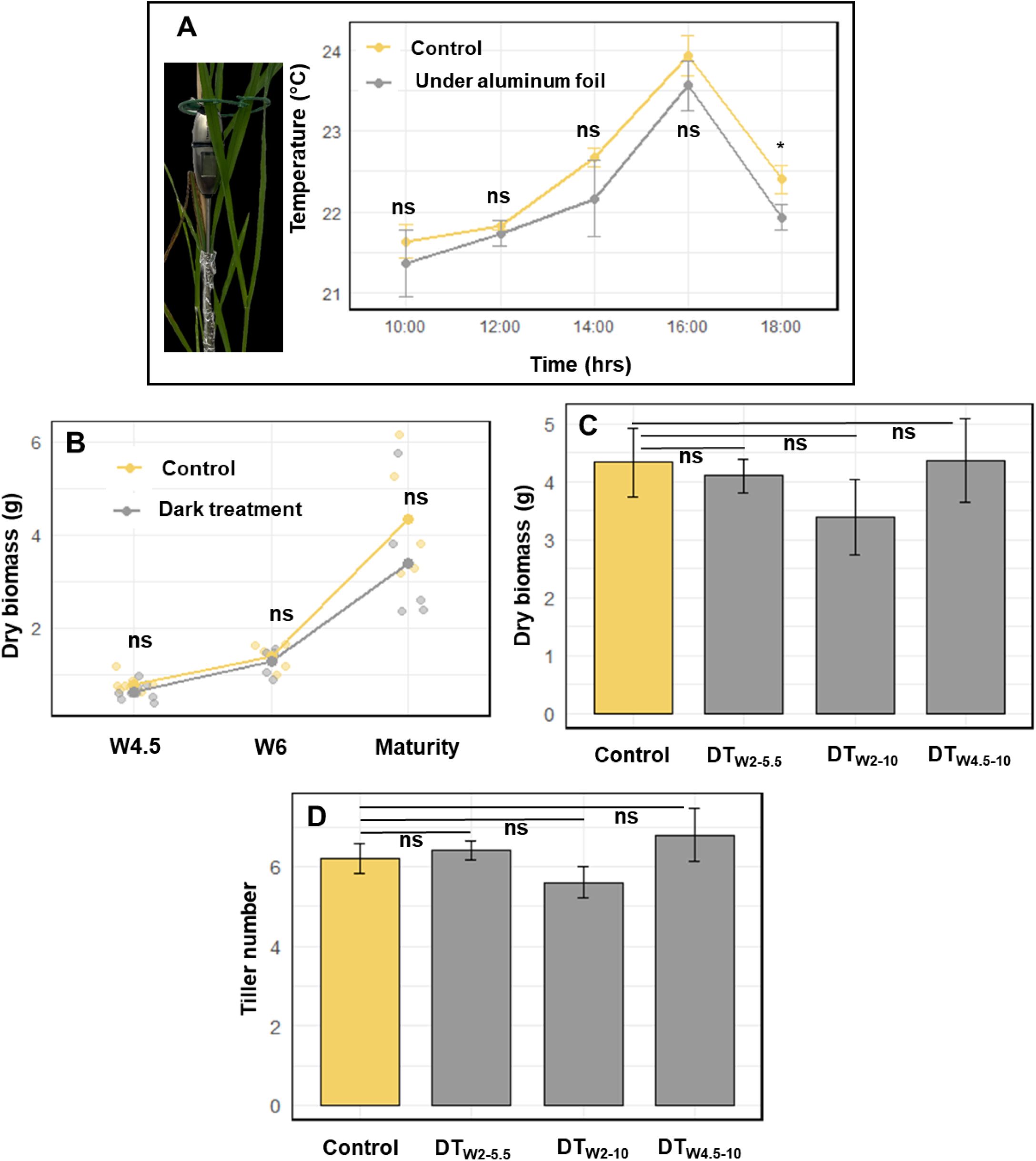
Effects of dark treatment on ambient temperature and plant growth. (A) Temperature measurements under aluminum foil covering the developing inflorescence compared to ambient temperature throughout the day. Comparison between control and dark treated plants for (B) Above ground dry biomass at W4.5, W6 and maturity. (C) Above ground dry biomass of whole plant. (D) Number of tillers per plant.

### 3.2 A nondestructive method to prevent pre-anthesis embryonic inflorescence greening (PEIG) using the dark treatment

We developed a technique to inhibit PEIG by covering the leaf sheaths surrounding the immature inflorescence with aluminum foil strips (Fig. 2b & S1), as described in detail in the materials and methods section. This dark treatment (DT) was applied to three distinct experimental groups (Fig. 2a). In the DT_W2-10_ group, the dark treatment began at developmental stage W2 and maintained until W10, marking the completion of inflorescence development until anthesis. In the DT_W2-5.5_ group, the treatment was applied from W2 to W5.5, approximately the midpoint of inflorescence development in barley; after W5.5, the covers were removed and the plants were exposed to normal greenhouse light conditions. In the DT_W4.5-10_ group, the treatment was applied from W4.5 to W10. A control group was continuously exposed to light throughout development. Meristems were dissected from the DT_W2-10_ and control groups across various developmental stages (W2-W10). From as early as stage W3-3.5, immature spikes from DT plants appeared pale and lacked visible chlorophyll compared to those of the control plants. This pale phenotype persisted throughout later stages of inflorescence development (W4+). Similarly, developing anthers from DT inflorescences exhibited no visible chlorophyll accumulation (Fig. 2b). These pale DT spikes also showed hardly any chlorophyll auto fluorescence, indicating a clear decline of chlorophyll (Fig. 2c). Quantification of chlorophyll pigments at stages W4.5 and W6 confirmed a significant reduction in chlorophyll (a + b), chlorophyll a and chlorophyll b levels in DT spikes at both stages (Fig. 2d-f).

To test whether the aluminum cover also affects the local temperature around and below the cover, the temperature beneath the foil was measured using a thermoprobe placed below the cover at multiple time points during the day (Fig. 3a). No significant temperature differences were observed between dark-treated and control plants, indicating that the treatment did not alter the local temperature around the developing inflorescence (Fig. 3a). In addition, agronomic traits, such as total tiller number at maturity, and dry biomass at different developmental stages and at maturity, were recorded for all treatment groups. The dry biomass did not change between DT and control plants across different developmental stages (Fig. 3b). In addition, for different treatment groups at maturity the dry biomass remained unchanged when compared to control (Fig. 3c). Furthermore, no significant differences were detected in the tiller number between the dark-treated and control plants (Fig. 3d), confirming that the DT specifically inhibited spike greening without affecting overall plant growth or development. Taken together, this study establishes a simple, non-destructive, and effective method to inhibit greening in immature inflorescences by preventing light exposure using a simple aluminum-cover technique.

### 3.3 Inhibition of PEIG impedes floral survival

To further evaluate whether the inhibition of PEIG had any influence on reproductive development; we analyzed the effect of the DT on spikelet initiation and survival dynamics. Morphological inspection of mature spikes revealed clear phenotypic differences between treatments. Spikes from DT_W2–10_ and DT_W4.5–10_ plants at maturity appeared noticeably smaller, weaker, and with very few grains compared to control spikes, whereas those from DT_W2–5.5_ plants could recover very efficiently and therefore appeared similar to controls (Fig. 4a). To quantify these visual differences, the final spikelet number was recorded at maturity. While DT_W2–10_ and DT_W4.5–10_ groups exhibited a significant reduction in final spikelet number (∼5-6 spikelets) compared with control plants, the earlier DT_W2–5.5_ treatment showed no significant difference in spikelet number relative to the control (Fig. 4c), indicating that light exposure during the later developmental period, i.e. W4.5-10, is essential for floral success.

**Figure 4:**
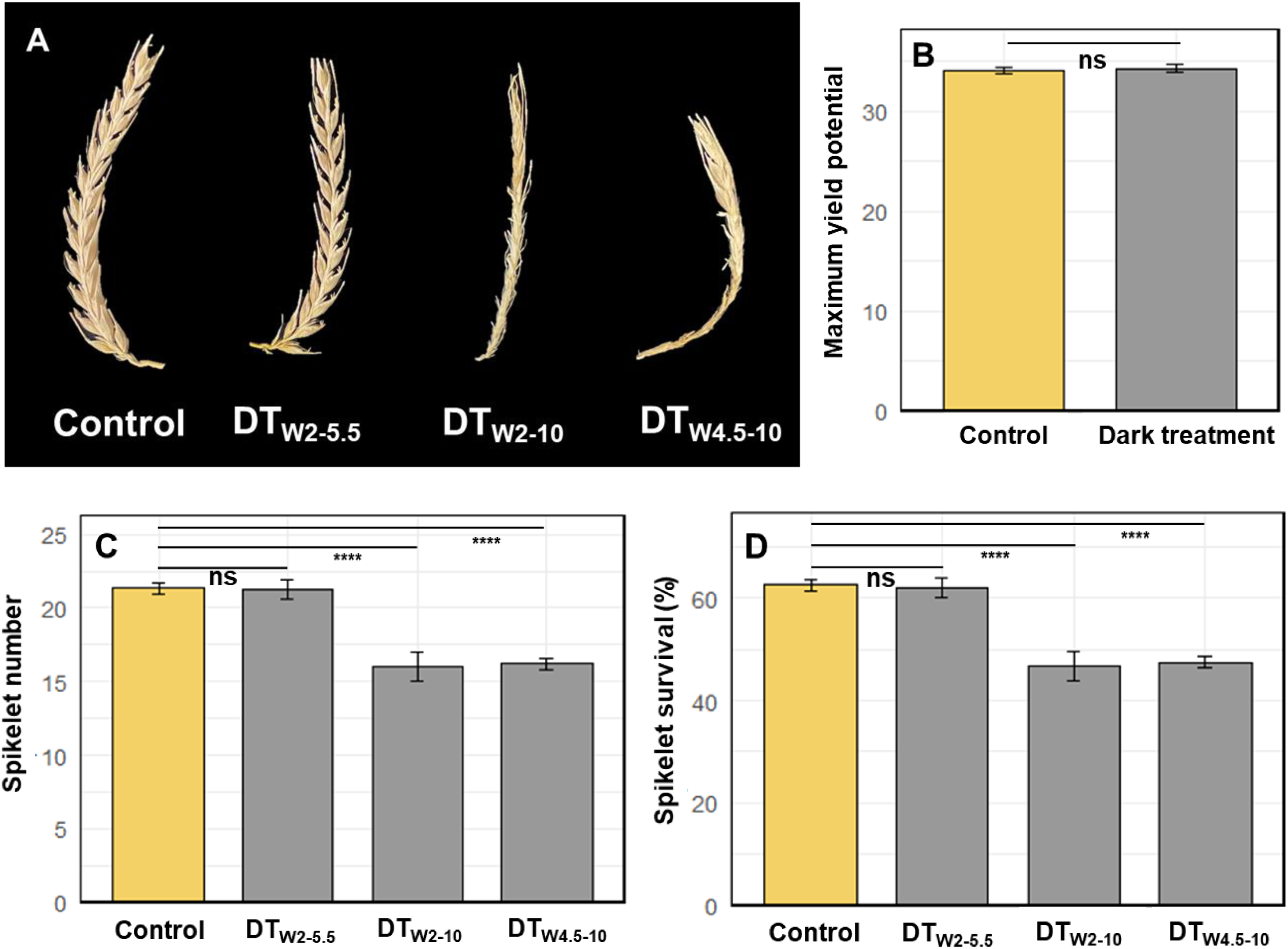
Effects of dark treatment on spike development. (A) Images of mature spikes from different treatment groups. (B) Maximum yield potential (MYP) of control and dark-treated spikes at W4.5. Comparison between treatment groups for (C) Spikelet number per spike, (D) Spikelet survival percentage per spike.

Further, the maximum yield potential (MYP), defined as the total number of spikelet primordia initiated per inflorescence, was determined for immature inflorescences collected from both dark-treated and control plants at the W4.5–5 developmental stage. This stage marks the period when spikelet initiation reaches its peak in barley, making it suitable for assessing potential developmental alterations. Importantly, the MYP did not differ significantly between any dark-treated and control spikes (Fig. 4b), proving that the number and rate of spikelet primordia initiation is independent of the greening process in the embryonic spike.

Spikelet survival was then calculated as the proportion of spikelets that survived at heading relative to the total number initiated. While DT_W2–10_ and DT_W4.5–10_ treatments showed distinctly increased spikelet mortality compared to controls, the DT_W2–5.5_ treatment exhibited spikelet survival rates almost identical to those of the control plants (Fig. 4d). The reduction in the spikelet number in the plants of DT_W2-10_ and DT_W4.5-10_ groups was therefore confirmed to be due to increased spikelet primordia degeneration. Taken together, these findings indicate that the prolonged absence of light and consequent inhibition of chlorophyll accumulation in developing inflorescences negatively affects floral fate during the later stages of spike maturation. Overall, these results demonstrate that while spikelet initiation is unaffected by the inhibition of greening, sustained dark treatment during later inflorescence development compromises spikelet survival, leading to a reduction in the final spikelet number.

### 3.4 Natural variation in PEIG affects spikelet survival

To investigate the effect of variable levels of PEIG among different barley genotypes, a set of 20 diverse six-rowed barley genotypes were selected from a broader diversity panel (Table S1). The selection was based on the extent of spikelet survival using the already published information (Huang et al., 2023), with the top 10 genotypes showing the highest spikelet survival and the bottom 10 genotypes exhibiting the lowest spikelet survival. This selection strategy was guided by previous observations in our study suggesting a possible relationship between PEIG and spikelet survival.

Chlorophyll content in immature spikes at the W4.5-5.5 developmental stage was quantified to assess the degree of greening across genotypes. A significant positive correlation was observed between spikelet survival and chlorophyll content, including total chlorophyll (a+b), chlorophyll a, and chlorophyll b (Fig. 5a-c). The chlorophyll a and a+b accounted for ∼60% (*r^2^* = 0.62 & 0.6, *p*<0.001) of the spikelet survival while chlorophyll b content accounted for ∼51% (*r^2^* = 0.51, *p*<0.001). Genotypes with higher chlorophyll content in their immature inflorescences, particularly within the apical region of the spike, e.g. HOR_11454, HOR_2503 and HOR_8446, exhibited markedly greater spikelet survival. In contrast, genotypes with reduced greening in both the whole spike and especially the apical portion, e.g. HOR_10560, HOR_11176 and HOR_15423, displayed a higher incidence of PTD (Fig 5d). These results demonstrate that the extent of PEIG, especially in the apical region, is closely linked to the degree of PTD that a spike undergoes, thereby influencing overall spikelet survival.

**Figure 5:**
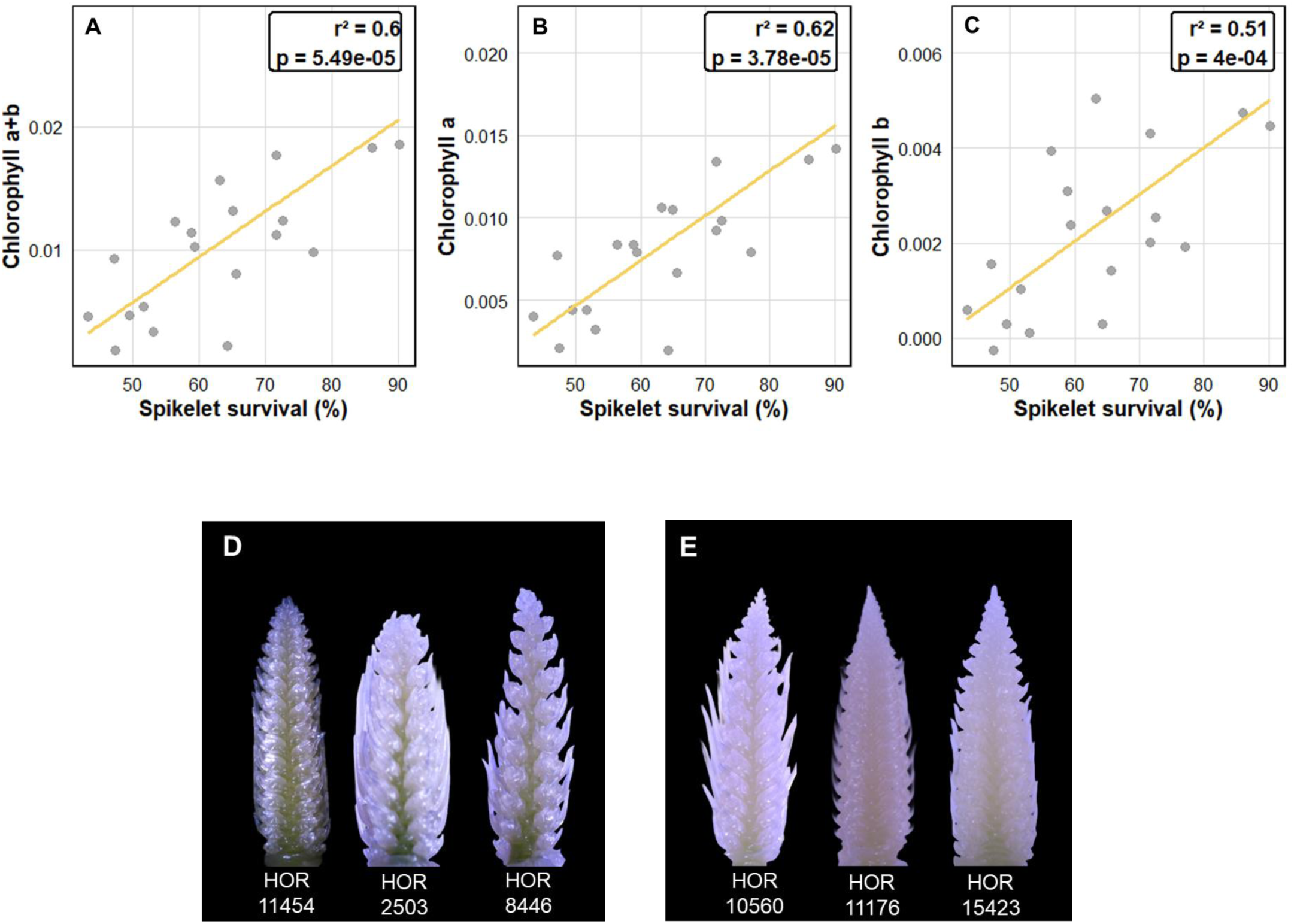
Relationship between chlorophyll content and spikelet survival. Correlation of spikelet survival percentage with: (A) Chlorophyll (a+b), (B) Chlorophyll a and (C) Chlorophyll b. Representative spike images at W4.5 of (D) High PEIG group and (E) low PEIG group

### 3.5 Influence of dark treatment on spikelet fertility

As already described in Results 3.1 and 3.2, developing anthers normally exhibit visible greening from very early stages of inflorescence development, indicating the onset of chlorophyll accumulation within these tissues. When greening was inhibited by applying the DT to the entire developing spike, including anthers, the anthers appeared pale and completely devoid of visible chlorophyll pigments (Fig. 2b).

In line with the pale anther phenotype, floral fertility was significantly reduced in the DT_W2-10_ and DT_W4.5-10_ groups compared with the control. In contrast, the DT_W2-5.5_ treatment showed no significant difference in the spike fertility relative to the control spikes (Fig. 6a). This trend closely resembled the observations of spikelet survival reported in the earlier results.

**Figure 6:**
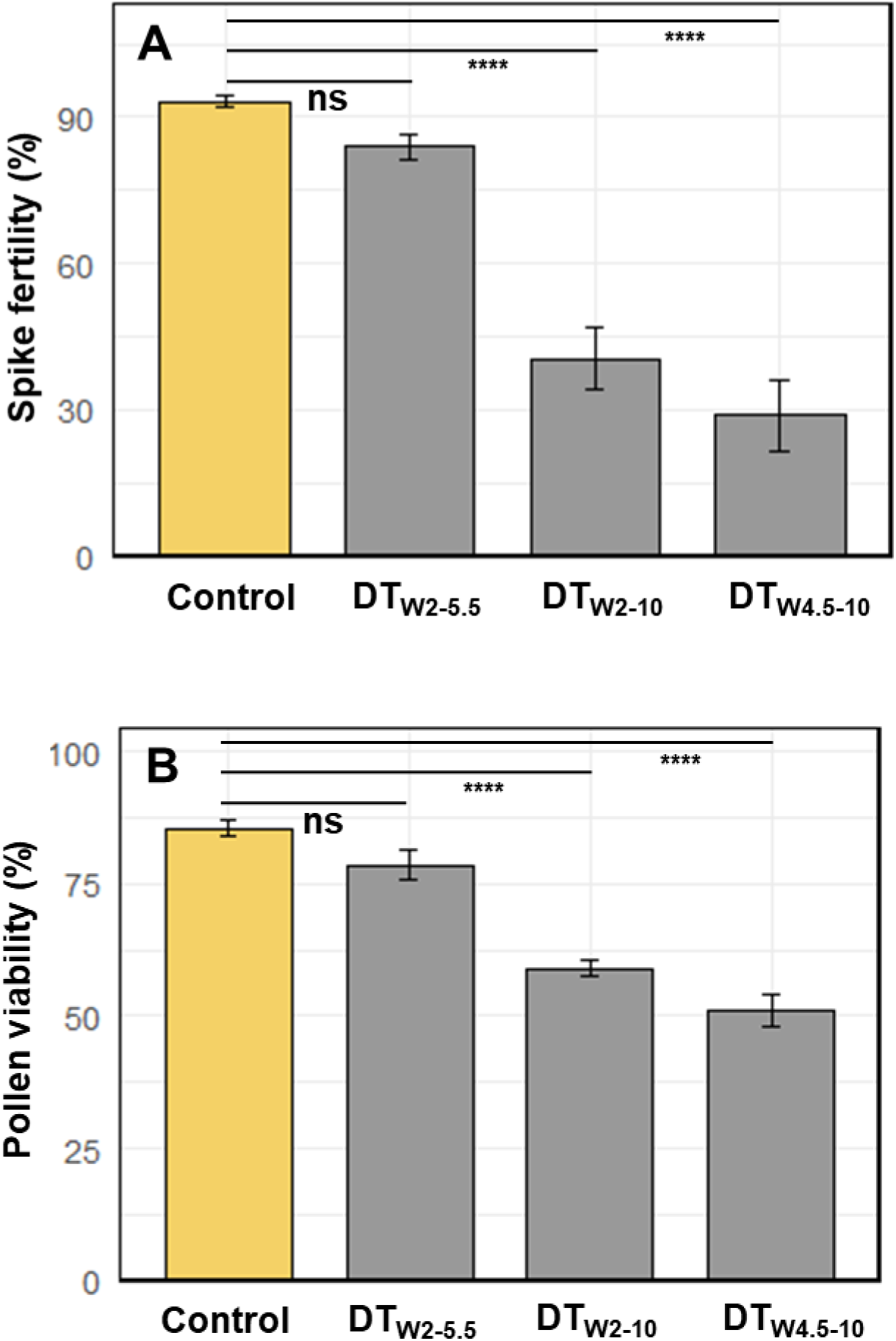
Effects of dark treatment on spike fertility. Comparison between different treatment groups for (A) Spike fertility percentage and (B) Pollen viability percentage

To further investigate the cause of reduced fertility in the DT plants, pollen quality was examined using impedance flow cytometry. As expected, both DT_W2-10_ and DT_W4.5-10_ spikes displayed a significant reduction in pollen viability compared with the control, while DT_W2-5.5_ spikes maintained normal pollen viability (Fig. 6b). These results demonstrate that the suppression of light-dependent greening in developing anthers directly compromises pollen development, leading to impaired fertility. Collectively, these findings indicate a strong association between anther greening and spikelet fertility, suggesting that chlorophyll biosynthesis or light perception within the anther tissues plays a crucial role in maintaining pollen viability and overall reproductive success in barley.

## 4. Discussion

Our study provides new insight into the developmental importance of light dependent chlorophyll accumulation during early inflorescence development in barley. Here, we demonstrate that (i) the PEIG begins at the earliest reproductive stage and follows a defined spatial patterning, (ii) nondestructive inhibition of PEIG through localized DT disrupts spikelet survival and pollen viability, and (iii) natural variation in PEIG across genotypes predicts differences in PTD. Together, these findings highlight a previously underappreciated role of localized photosynthetic competence within developing inflorescence and anthers in shaping reproductive success in cereals.

### 4.1 Timing and spatial progression of chlorophyll accumulation in immature inflorescences

Chlorophyll biosynthesis begins around W1.5, which is when the shoot apex shifts to reproductive development (Waddington et al., 1983). According to the analyses of chlorophyll autofluorescence, the central developing rachis was the primary site of early greening, which subsequently spread to spikelet primordia and floral organs. This progression closely parallels cellular differentiation and tissue expansion within the inflorescence (Kirby & Appleyard, 1984), suggesting that chlorophyll biogenesis is developmentally coordinated with the spike’s architectural establishment. Leaves are traditionally considered to be primary photosynthetic organs in higher plants and major carbon sources (Van Camp, 2005; White et al., 2016), whereas reproductive structures are commonly considered carbon sinks that depend largely on leaf derived photoassimilates (Van Camp, 2005; White et al., 2016). However, several evidences indicate that reproductive organs can make significant contribution of photoassimilates (Aschan et al., 2003; Brazel et al., 2019; Raven et al., 2015), particularly during the later stages of development (Noodén et al., 2001). Our findings may extend this concept by demonstrating that immature barley inflorescence accumulate chlorophyll very early during development, even under the limited light that penetrates between the leaf sheaths to reach the developing inflorescence. This early biogenesis raises the possibility that developing spikes possess at least a limited potential for localized photosynthesis or chlorophyll mediated stress signaling (M. Li & Kim, 2022; Trentmann et al., 2020). However, direct assessment of photosynthetic machinery and its competence in these tissues will be required in order to better understand the exact role of PEIG in producing photoassimilates.

### 4.2 Light signal mediated inflorescence greening

In our study, we demonstrate that the light signal was directly perceived by the developing and sheathed inflorescence. The aluminum foil mediated DT method developed here provides a simple and non-invasive method to block light perception and further chlorophyll accumulation in the developing spikes without affecting whole plant growth or local temperature. It has been previously reported that the altered temperature (Hu et al., 2021; Jagadish, 2020; Prasad & Djanaguiraman, 2014; Saini & Aspinall, 1982) or altered development (Willey & Holliday, 1971) can significantly disrupt the inflorescence development and increase sterility. Therefore, our results, which display no significant difference in the ambient temperature upon aluminum foil cover and in the dry biomass or the tiller number, ensure that the observed phenotypes were a direct consequence of suppressed chlorophyll biosynthesis in the developing spike. The pale phenotype of both developing floral structures and anthers from early stages (W3-3.5) onward, together with significant reductions in chlorophyll a, b, and total chlorophyll levels, underscores the essential role of light perception in promoting plastid differentiation and chlorophyll accumulation. These findings are consistent with the well-established requirement of light-regulated transcriptional and plastid developmental pathways for chlorophyll biosynthesis (Chen et al., 2004; Jarvis & López-Juez, 2013; Pogson & Albrecht, 2011; Yuan et al., 2017)). Consequently, the absence of any light disrupts these coordinated biogenetic processes in developing inflorescence tissues, leading to a persistent chlorophyll-deficient phenotype. In contrast, the initiation of chlorophyll biosynthesis in control spikes indicates that some amount of light can penetrate through the surrounding leaf sheath layers to reach the immature inflorescence, providing a critical cue for chlorophyll accumulation. Although most of the blue and red wavelengths of the visible light spectrum are absorbed by the surrounding leaf sheath tissues, unabsorbed green light may still reach the developing inflorescence. While often underappreciated as a signaling cue, several studies have shown that monochromatic green light can trigger specific developmental responses in plants, such as hypocotyl elongation in Arabidopsis, accumulation of geranylgeranylated chlorophylls in barley, and altered developmental rates in wheat (Battle et al., 2020; Kasajima et al., 2009; Materová et al., 2017; Smith et al., 2017). Previous work has also demonstrated that light can be transmitted through intercellular air spaces or vascular tissues, providing sufficient illumination to activate the light-dependent gene networks necessary for early chloroplast development (Nawkar et al., 2023; Sun et al., 2003, 2005).

### 4.3 PEIG as a physiological determinant of overall reproductive fitness in cereals

Selectively inhibiting the light revealed that prolonged darkening (DT_W2-10_ or DT_W4.5-10_) drastically reduced final spikelet number, whereas early short-term inhibition (DT_W2–5.5_) allowed full recovery. Interestingly, maximum primordia number remained unchanged. Therefore, the reductions in final spikelet number were attributable to increased PTD during later developmental stages (Huang et al., 2023; Shanmugaraj et al., 2023; Thirulogachandar & Schnurbusch, 2021). PTD is a major determinant of grain number in barley (Huang et al., 2023; Kamal, Muqaddasi, Zhao, et al., 2022; Thirulogachandar & Schnurbusch, 2021), and is strongly influenced by local assimilate supply and developmental stability (Shanmugaraj et al., 2023; Sreenivasulu & Schnurbusch, 2012). The pronounced effects of light deprivation suggest that developing spikes may rely on chloroplast-derived cues to support rapid cellular differentiation and establish proper tissue identity. Chloroplasts play a central role in guiding cell fate and tissue differentiation, ensuring that cells develop appropriately for their location and function (Guo et al., 2016; Liebers et al., 2022; Sierra et al., 2023). In cereal spikes, early chloroplast development likely provides signals that coordinate spikelet survival, vascular differentiation, and overall inflorescence architecture, in addition to later supplying photoassimilates once tissues are fully differentiated. This is consistent with evidence that cereal inflorescences can act as both sinks and modest photosynthetic sources at later stages of development (AuBuchon-Elder et al., 2020; Bort et al., 1994; Maydup et al., 2012; Sanchez-Bragado et al., 2014; Tambussi et al., 2005). Natural variation among 20 barley genotypes further supported this functional link: chlorophyll content in immature spikes, explained 50–62% of variation in spikelet survival, and genotypes with low PEIG showed increased tip degeneration, aligning with known genotypic differences in PTD or spikelet survival in barley (Kamal, Muqaddasi, & Schnurbusch, 2022; Kamal, Muqaddasi, Zhao, et al., 2022). PEIG therefore emerges as a quantitative trait influencing reproductive fitness and offers potential breeding value, especially given the identified QTLs for spikelet survival and the association of the candidate gene with PEIG (Huang et al., 2023). Notably, PEIG is not restricted to barley but is a characteristic feature of several major cereals including wheat and rye suggesting that early inflorescence greening represents a conserved developmental strategy that can be exploited as an important trait for crop improvement across Triticeae species. Beyond spikelets, we also found that anther greening initiated early during microsporogenesis. As reported earlier, light signal influences spike fertility (Dreccer et al., 2022; González et al., 2003; Ugarte et al., 2010). Similarly, also in our study, dark-treated developing anthers remained pale and produced largely non-viable pollen, which in turn hampers the overall spike fertility. This indicates that light-dependent chloroplast maturation may support the tapetum function, lipid biosynthesis, and microspore metabolism (Ariizumi & Toriyama, 2011; Zhu et al., 2020). Similar sterility phenotypes are observed in mutants deficient in chloroplast development (Zhu et al., 2020). Although, here we propose that the lack of light hampers the pollen viability and fertility, the possible effect from the female reproductive organs on spikelet sterility needs to be investigated. Future experiments incorporating recovery assays, or direct morphological analyses of female reproductive organ development will be crucial to fully elucidate the contribution of female defects to spikelet sterility. Collectively, these results establish that light perception and greening in the developing inflorescence of barley and other cereals are not passive traits but integral processes that enhance spikelet retention and pollen fertility, thereby underpinning overall reproductive fitness.

## Acknowledgments

We greatfully thank K.Wolf, C.Trautewig and E. Weiss for their excellent technical support; K.G. Koch and her team for greenhouse plant care; D. Kumar for helping with pollen viability experiments, C. Waesch for critically reviewing previous versions of the manuscript, and all the members of the plant architecture group at IPK for fruitful discussions. Infrastructure and financial support for this project was received from the IPK core budget as part of a seed-funding project for establishing joint scientific links between CEPLAS (Cluster of Excellence on Plant Sciences) and IPK. CEPLAS is funded by the Deutsche Forschungsgemeinschaft under Germany’s Excellence Strategy EXC-2048/1 under project ID 390686111.

## Author contributions

T.S., Y.H., and A.P.M.W. conceived the project, were involved in designing experiments and supervised the study. S.T.B. designed and conducted all the experiments, collected the phenotypic data, performed the data analyses, designed the figures, and wrote the first draft of the manuscript; Y.H. provided phenotypic data for 350 barley lines. All authors have read and approved the manuscript.

---

**Figure S1**: **Dark-treatment procedure used to prevent light exposure to the developing inflorescence at ∼W4.5.** (A) Identification of the position of the developing spike. (B) Wrapping aluminum foil around the leaf sheaths surrounding the internode at the spike position. (C) Ensuring coverage of the leaf sheaths approximately 2–3 cm above and below the spike to maintain complete darkness. (a) Distance between the internode bearing developing spike and ground, (b) Internode bearing the developing inflorescence, (c) Total area covered by aluminum foil.

**Figure S2: A systematic illustration of the procedure used to measure temperature beneath the aluminum-foil cover.** (A) Identification of the position of the developing spike. (B) Placement of the thermometer such that the probe tip sits at the internode where the spike is developing. (C) Wrapping the aluminum foil around the leaf sheath and thermometer probe.(D) Providing a support stick to stabilize the thermometer. (a) Long-probe digital thermometer; (b) distance between the internode bearing the developing spike and the ground; (c) internode bearing the developing spike.

## References

Ariizumi, T., & Toriyama, K. (2011). Genetic regulation of sporopollenin synthesis and pollen exine development. Annual Review of Plant Biology, 62, 437–460

Arisnabarreta, S., & Miralles, D. J. (2008). Critical period for grain number establishment of near isogenic lines of two- and six-rowed barley. Field Crops Research, 107(3), 196–202.

Aschan, G., & Pfanz, H. (2003). Non-foliar photosynthesis – a strategy of additional carbon acquisition. Flora - Morphology, Distribution, Functional Ecology of Plants, 198(2), 81–97.

AuBuchon-Elder, T., Coneva, V., Goad, D. M., Jenkins, L. M., Yu, Y., Allen, D. K., & Kellogg, E. A. (2020). Sterile Spikelets Contribute to Yield in Sorghum and Related Grasses. The Plant Cell, 32(11), 3500–3518.

Battle, M. W., Vegliani, F., & Jones, M. A. (2020). Shades of green: untying the knots of green photoperception. Journal of Experimental Botany, 71(19), 5764–5770.

Bort, J., Febrero, A., Amaro, T., & Araus, J. L. (1994). Role of awns in ear water-use efficiency and grain weight in barley Plant physiology Role of awns in ear water-use efficiency and grain weight in barley. 14(2), 133–139.

Brazel, A. J., & Ó’Maoileídigh, D. S. (2019). Photosynthetic activity of reproductive organs. Journal of Experimental Botany, 70(6), 1737–1754.

Cackett, L., Luginbuehl, L. H., Schreier, T. B., Lopez-Juez, E., & Hibberd, J. M. (2022). Chloroplast development in green plant tissues: the interplay between light, hormone, and transcriptional regulation. New Phytologist, 233(5), 2000–2016.

Chen, M., Chory, J., & Fankhauser, C. (2004). Light signal transduction in higher plants. Annual Review of Genetics, 38(Volume 38, 2004), 87–117.

Dong, B., Yang, H., Liu, H., Qiao, Y., Zhang, M., Wang, Y., Xie, Z., & Liu, M. (2019). Effects of shading stress on grain number, yield, and photosynthesis during early reproductive growth in wheat. Crop Science, 59(1), 363–378.

Dreccer, M. F., Zwart, A. B., Schmidt, R. C., Condon, A. G., Awasi, M. A., Grant, T. J., Galle, A., Bourot, S., & Frohberg, C. (2022). Wheat yield potential can be maximized by increasing red to far-red light conditions at critical developmental stages. Plant Cell and Environment, 45(9), 2652–2670.

Gebbing, T., & Schnyder, H. (2001). 13C Labeling kinetics of sucrose in glumes indicates significant refixation of respiratory CO2 in the wheat ear. Australian Journal of Plant Physiology, 28(10), 1047–1053.

González, F. G., Slafer, G. A., & Miralles, D. J. (2003). Floret development and spike growth as affected by photoperiod during stem elongation in wheat. Field Crops Research, 81(1), 29–38.

González, F. G., Slafer, G. A., & Miralles, D. J. (2005). Floret development and survival in wheat plants exposed to contrasting photoperiod and radiation environments during stem elongation. Functional Plant Biology : FPB, 32(3), 189–197.

Guo, H., Feng, P., Chi, W., Sun, X., Xu, X., Li, Y., Ren, D., Lu, C., David Rochaix, J., Leister, D., & Zhang, L. (2016). Plastid-nucleus communication involves calcium-modulated MAPK signalling. Nature Communications, 7, 12173.

Huang, Y., Kamal, R., Shanmugaraj, N., Rutten, T., Thirulogachandar, V., Zhao, S., Hoffie, I., Hensel, G., Rajaraman, J., Moya, Y. A. T., Hajirezaei, M. R., Himmelbach, A., Poursarebani, N., Lundqvist, U., Kumlehn, J., Stein, N., von Wirén, N., Mascher, M., Melzer, M., & Schnurbusch, T. (2023). A molecular framework for grain number determination in barley. Science Advances, 9(9).

Hu, Q., Wang, W., Lu, Q., Huang, J., Peng, S., & Cui, K. (2021). Abnormal anther development leads to lower spikelet fertility in rice (Oryza sativa L.) under high temperature during the panicle initiation stage. BMC Plant Biology 2021 21:1, 21(1), 428-.

Imaizumi, N., Usuda, H., Nakamoto, H., & Ishihara, K. (1990). Changes in the Rate of Photosynthesis during Grain Filling and the Enzymatic Activities Associated with the Photosynthetic Carbon Metabolism in Rice Panicles. Plant and Cell Physiology, 31(6), 835–844.

Jagadish, S. V. K. (2020). Heat stress during flowering in cereals – effects and adaptation strategies. New Phytologist, 226(6), 1567–1572.

Jarvis, P., & López-Juez, E. (2013). Biogenesis and homeostasis of chloroplasts and other plastids. Nature Reviews Molecular Cell Biology, 14(12), 787–802.

Kamal, R., Muqaddasi, Q. H., & Schnurbusch, T. (2022). Genetic association of spikelet abortion with spike, grain, and shoot traits in highly-diverse six-rowed barley. Frontiers in Plant Science, 13, 1015609.

Kamal, R., Muqaddasi, Q. H., Zhao, Y., & Schnurbusch, T. (2022). Spikelet abortion in six-rowed barley is mainly influenced by final spikelet number, with potential spikelet number acting as a suppressor trait. Journal of Experimental Botany, 73(7), 2005–2020.

Kasajima, S. ya, Inoue, N., & Mahmud, R. (2009). Response spectrum for green light-induced acceleration of heading in wheat cv. Norin 61. Plant Production Science, 12(1), 54–57.

Koppolu, R., & Schnurbusch, T. (2019). Developmental pathways for shaping spike inflorescence architecture in barley and wheat. Journal of Integrative Plant Biology, 61(3), 278–295.

Liebers, M., Cozzi, C., Uecker, F., Chambon, L., Blanvillain, R., & Pfannschmidt, T. (2022). Biogenic signals from plastids and their role in chloroplast development. Journal of Experimental Botany, 73(21), 7105–7125.

Li, M., & Kim, C. (2022). Chloroplast ROS and stress signaling. Plant Communications, 3(1), 100264.

Li, X., Wang, H., Li, H., Zhang, L., Teng, N., Lin, Q., Wang, J., Kuang, T., Li, Z., Li, B., Zhang, A., & Lin, J. (2006). Awns play a dominant role in carbohydrate production during the grain-filling stages in wheat (Triticum aestivum). Physiologia Plantarum, 127(4), 701–709.

Masanori, T., Ichiro, T., Akihito, K., & Koh-Ichiro, A. (2001). Initiation and development of spikelets and florets in wheat as influenced by shading and nitrogen supply at the spikelet phase. Plant Production Science, 4(4), 283–290.

Materová, Z., Sobotka, R., Zdvihalová, B., Oravec, M., Nezval, J., Karlický, V., Vrábl, D., Štroch, M., & Špunda, V. (2017). Monochromatic green light induces an aberrant accumulation of geranylgeranyled chlorophylls in plants. Plant Physiology and Biochemistry, 116, 48–56.

Maydup, M. L., Antonietta, M., Guiamet, J. J., & Tambussi, E. A. (2012). The contribution of green parts of the ear to grain filling in old and modern cultivars of bread wheat (Triticum aestivum L.): Evidence for genetic gains over the past century. Field Crops Research, 134, 208–215.

Mckenzie, H. (2011). ADVERSE INFLUENCE OF AWNS ON YIELD OF WHEAT. 10.4141/Cjps72-010, 52(1), 81–87.

Nawkar, G. M., Legris, M., Goyal, A., Schmid-Siegert, E., Fleury, J., Mucciolo, A., De Bellis, D., Trevisan, M., Schueler, A., & Fankhauser, C. (2023). Air channels create a directional light signal to regulate hypocotyl phototropism. Science, 382(6673), 935–940.

Noodén, L., Botany, J. P.-J. of E., & 2001, undefined. (n.d.). Correlative controls of senescence and plant death in Arabidopsis thaliana (Brassicaceae). Journal of Experimental Botany.

Parrado, J. D., Savin, R., & Slafer, G. A. (2025). Floret development and fertility of barley as affected by Photoperiod-H1 under contrasting photoperiods and PHYTOCHROME C backgrounds. Journal of Experimental Botany, 76(6), 1691–1703.

Parrado, J. D., Slafer, G. A., & Savin, R. (2025). Diverse alleles of Photoperiod-H1 directly and indirectly affect barley yield-related traits under contrasting photoperiods and PHYTOCHROME C backgrounds. Journal of Experimental Botany, 76(6), 1678–1690.

Pogson, B. J., & Albrecht, V. (2011). Genetic Dissection of Chloroplast Biogenesis and Development: An Overview. Plant Physiology, 155(4), 1545–1551.

Pogson, B. J., Ganguly, D., & Albrecht-Borth, V. (2015). Insights into chloroplast biogenesis and development. Biochimica et Biophysica Acta (BBA) - Bioenergetics, 1847(9), 1017–1024.

Porra, R. J., Thompson, W. A., & Kriedemann, P. E. (1989). Determination of accurate extinction coefficients and simultaneous equations for assaying chlorophylls a and b extracted with four different solvents: verification of the concentration of chlorophyll standards by atomic absorption spectroscopy. Biochimica et Biophysica Acta (BBA) - Bioenergetics, 975(3), 384–394.

Prasad, P. V. V., & Djanaguiraman, M. (2014). Response of floret fertility and individual grain weight of wheat to high temperature stress: sensitive stages and thresholds for temperature and duration. Functional Plant Biology, 41(12), 1261–1269.

Raven, J. A., & Griffiths, H. (2015). Photosynthesis in reproductive structures: costs and benefits. Journal of Experimental Botany, 66(7), 1699–1705.

Saini, H. S., & Aspinall, D. (1982). Abnormal Sporogenesis in Wheat (Triticum aestivum L.) Induced by Short Periods of High Temperature. Annals of Botany, 49(6), 835–846.

Sanchez-Bragado, R., Molero, G., Reynolds, M. P., & Araus, J. L. (2014). Relative contribution of shoot and ear photosynthesis to grain filling in wheat under good agronomical conditions assessed by differential organ δ13C. Journal of Experimental Botany, 65(18), 5401–5413.

Sanchez-Bragado, R., Molero, G., Reynolds, M. P., & Araus, J. L. (2016). Photosynthetic contribution of the ear to grain filling in wheat: a comparison of different methodologies for evaluation. Journal of Experimental Botany, 67(9), 2787–2798.

Shanmugaraj, N., Rajaraman, J., Kale, S., Kamal, R., Huang, Y., Thirulogachandar, V., Garibay-Hernández, A., Budhagatapalli, N., Tandron Moya, Y. A., Hajirezaei, M. R., Rutten, T., Hensel, G., Melzer, M., Kumlehn, J., von Wirén, N., Mock, H.-P., & Schnurbusch, T. (2023). Multilayered regulation of developmentally programmed pre-anthesis tip degeneration of the barley inflorescence. The Plant Cell, 35(11), 3973–4001.

Sierra, J., Escobar-Tovar, L., & Leon, P. (2023). Plastids: diving into their diversity, their functions, and their role in plant development. Journal of Experimental Botany, 74(8), 2508–2526.

Slafer, G. A., Calderini, D. F., Miralles, D. J., & Dreccer, M. F. (1994). Preanthesis shading effects on the number of grains of three bread wheat cultivars of different potential number of grains. Field Crops Research, 36(1), 31–39.

Smith, H. L., Mcausland, L., & Murchie, E. H. (2017). Don’t ignore the green light: exploring diverse roles in plant processes. Journal of Experimental Botany, 68(9), 2099–2110.

Sreenivasulu, N., & Schnurbusch, T. (2012). A genetic playground for enhancing grain number in cereals. Trends in Plant Science, 17(2), 91–101.

Sun, Q., Yoda, K., & Suzuki, H. (2005). Internal axial light conduction in the stems and roots of herbaceous plants. Journal of Experimental Botany, 56(409), 191–203.

Sun, Q., Yoda, K., Suzuki, M., & Suzuki, H. (2003). Vascular tissue in the stem and roots of woody plants can conduct light. Journal of Experimental Botany, 54(387), 1627–1635.

Tambussi, E. A., Nogués, S., & Araus, J. L. (2005). Ear of durum wheat under water stress: water relations and photosynthetic metabolism. Planta 2005 221:3, 221(3), 446–458.

Thirulogachandar, V., & Schnurbusch, T. (2021). ‘Spikelet stop’ determines the maximum yield potential stage in barley. Journal of Experimental Botany, 72(22), 7743–7753.

Trentmann, O., Mühlhaus, T., Zimmer, D., Sommer, F., Schroda, M., Haferkamp, I., Keller, I., Pommerrenig, B., & Neuhaus, H. E. (2020). Identification of Chloroplast Envelope Proteins with Critical Importance for Cold Acclimation. Plant Physiology, 182(3), 1239–1255.

Ugarte, C. C., Trupkin, S. A., Ghiglione, H., Slafer, G., & Casal, J. J. (2010). Low red/far-red ratios delay spike and stem growth in wheat. Journal of Experimental Botany, 61(11), 3151–3162.

Van Camp, W. (2005). Yield enhancement genes: seeds for growth. Current Opinion in Biotechnology, 16(2), 147–153.

Waddington, S. R., Cartwright, P. M., & Wall, P. C. (1983a). A Quantitative Scale of Spike Initial and Pistil Development in Barley and Wheat. Annals of Botany, 51(1), 119–130.

Waddington, S. R., Cartwright, P. M., & Wall, P. C. (1983b). A Quantitative Scale of Spike Initial and Pistil Development in Barley and Wheat. Annals of Botany, 51(1), 119–130.

White, A. C., Rogers, A., Rees, M., & Osborne, C. P. (2016). How can we make plants grow faster? A source–sink perspective on growth rate. Journal of Experimental Botany, 67(1), 31–45.

Willey, R. W., & Holliday, R. (1971). Plant population and shading studies in barley. The Journal of Agricultural Science, 77(3), 445–452.

Yang, H., Dong, B., Wang, Y., Qiao, Y., Shi, C., Jin, L., & Liu, M. (2020). Photosynthetic base of reduced grain yield by shading stress during the early reproductive stage of two wheat cultivars. Scientific Reports 2020 10:1, 10(1), 14353-.

Yuan, M., Zhao, Y. Q., Zhang, Z. W., Chen, Y. E., Ding, C. B., & Yuan, S. (2017). Light Regulates Transcription of Chlorophyll Biosynthetic Genes During Chloroplast Biogenesis. Critical Reviews in Plant Sciences, 36(1), 35–54.

Zhu, T., Li, Z., An, X., Long, Y., Xue, X., Xie, K., Ma, B., Zhang, D., Guan, Y., Niu, C., Dong, Z., Hou, Q., Zhao, L., Wu, S., Li, J., Jin, W., & Wan, X. (2020). Normal Structure and Function of Endothecium Chloroplasts Maintained by ZmMs33-Mediated Lipid Biosynthesis in Tapetal Cells Are Critical for Anther Development in Maize. Molecular Plant, 13(11), 1624–1643.

