## Supplementary material for "Pre-anthesis light signaling of sheathed embryonic barley inflorescences defines floral fate": Fig. S1

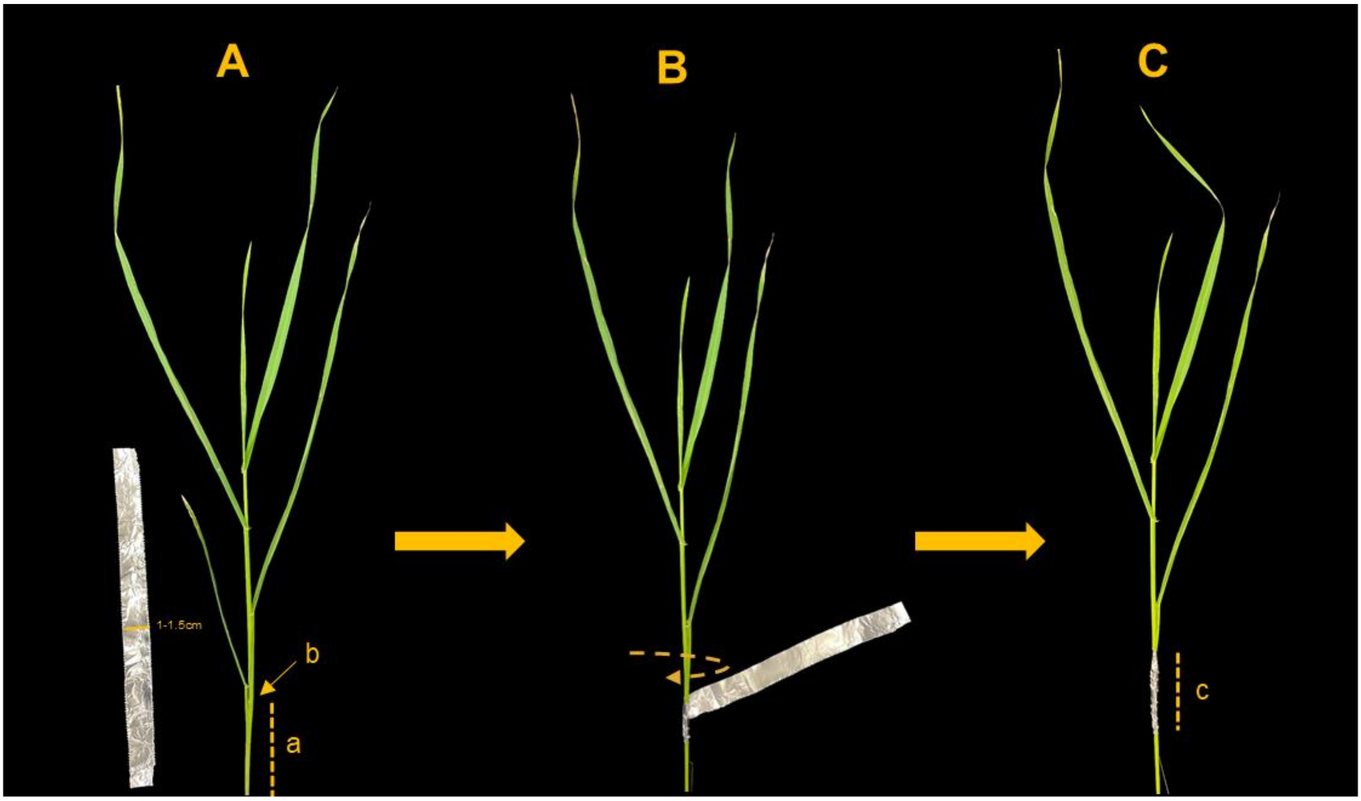


**Figure S1 - Dark-treatment procedure used to prevent light exposure to the developing inflorescence at ~W4.5.** (A) Identification of the position of the developing spike. (B) Wrapping aluminum foil around the leaf sheaths surrounding the internode at the spike position. (C) Ensuring coverage of the leaf sheaths approximately 2–3 cm above and below the spike to maintain complete darkness. (a) Distance between the internode bearing developing spike and ground, (b) Internode bearing the developing inflorescence, (c) Total area covered by aluminum foil.
