## Supplementary material for "Pre-anthesis light signaling of sheathed embryonic barley inflorescences defines floral fate": Fig. S2

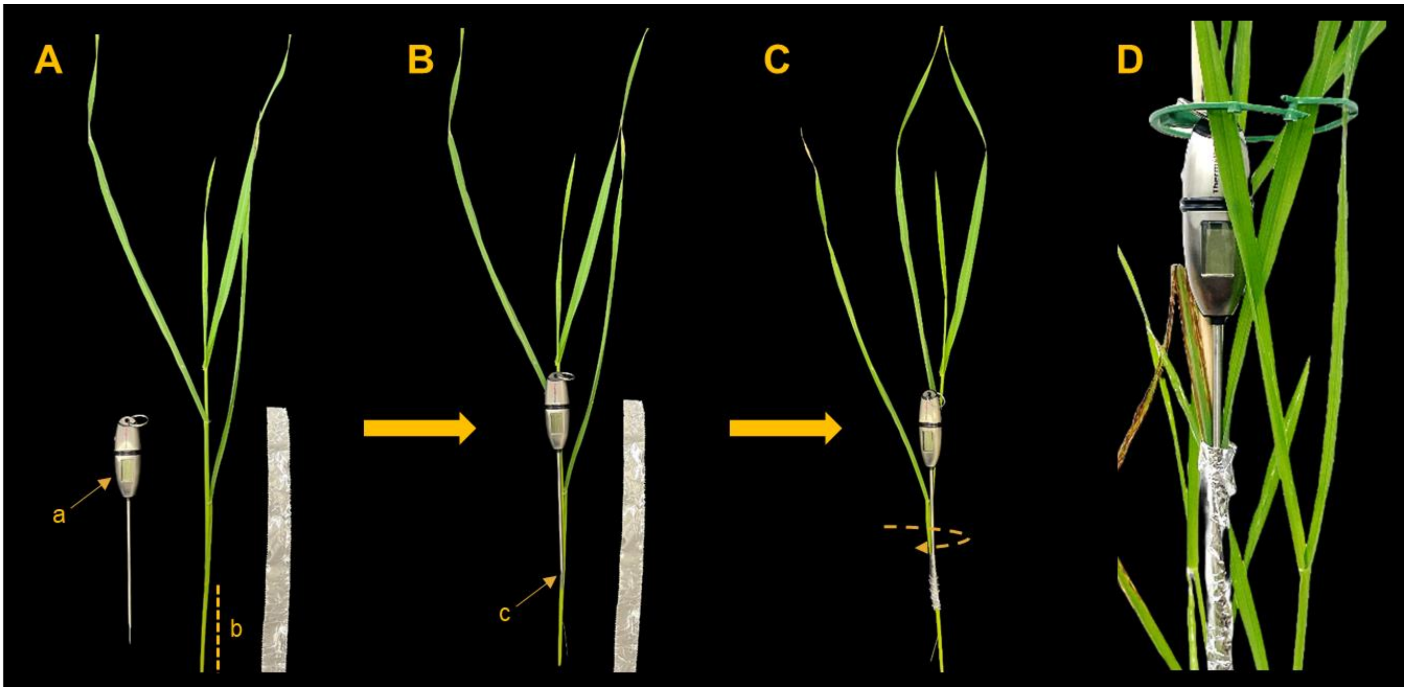


**Fig. S2: A step by step illustration of the procedure used to measure temperature beneath the aluminum-foil cover.** (A) Identification of the position of the developing spike. (B) Placement of the thermometer such that the probe tip sits at the internode where the spike is developing. (C) Wrapping the aluminum foil around the leaf sheath and thermometer probe. (D) Providing a support stick to stabilize the thermometer. (a) Long-probe digital thermometer; (b) distance between the internode bearing the developing spike and the ground; (c) internode bearing the developing spike.
